# panModule: detecting conserved modules in the variable regions of a pangenome graph

**DOI:** 10.1101/2021.12.06.471380

**Authors:** Adelme Bazin, Claudine Medigue, David Vallenet, Alexandra Calteau

## Abstract

The recent years have seen the rise of pangenomes as comparative genomic tools to better understand the evolution of gene content among microbial genomes in close phylogenetic groups such as species. While the core or persistent genome is often well-known as it includes essential or ubiquitous genes, the variable genome is usually less characterized and includes many genes with unknown functions even among the most studied organisms. It gathers important genes for strain adaptation that are acquired by horizontal gene transfer. Here, we introduce panModule, an original method to identify conserved modules in pangenome graphs built from thousands of microbial genomes. These modules correspond to synteny blocks composed of consecutive genes that are conserved in a subset of the compared strains. Identifying conserved modules can provide insights on genes involved in the same functional processes, and as such is a very helpful tool to facilitate the understanding of genomic regions with complex evolutionary histories. The panModule method was benchmarked on a curated dataset of conserved modules in *Escherichia coli* genomes. Its use was illustrated through a study of a high pathogenicity island in *Klebsiella pneumoniae* that allowed a better understanding of this region. panModule is freely available and accessible through the PPanGGOLiN software suite (https://github.com/labgem/PPanGGOLiN).

## 1 Introduction

Lately, the data deluge provided by NGS has given access to over a million of prokaryotic genome sequences in public data banks, as well as a wealth of genomes reconstructed from environmental data, such as metagenome assembled genomes (MAGs) or single-cell assembled genomes (SAGs). Consequently, many bacterial species of interest now have hundreds to several thousands of genomes publicly available. It represents a fantastic opportunity to understand the evolution of prokaryotic genomes, and more specifically to study gene flow within a given species. However, processing such a huge amount of data for comparative genomics analyses is becoming a real challenge and requires new bioinformatics solutions. In particular, many new methods rely on the conceptual framework of the pangenome, which corresponds to the entire gene repertoire of a taxonomic group [1]. Multiple methods have been developed to study pangenome structures and to perform comparative studies of hundreds to thousands of genomes (e.g. Roary [2], PIRATE [3], Panaroo [4], PPanGGOLiN [5], PanACoTA [6]). A pangenome can be described by two components: the core genome, which contains genes shared by all (or almost all) individuals, and the variable genome, which gathers all other genes.

Studying the variable part of the pangenome is of major interest since it gathers genes of importance for adaptation of the strains, and particularly genes that have been acquired by Horizontal gene transfer (HGT). Indeed, as described for many years, HGT is a significant source of gene novelty [7] and is a major driver of genome evolution in bacterial species providing and maintaining diversity at the population level [8, 9]. This horizontal gene flow occurs through different well-known mechanisms involving mobile genetic elements and can spread genes between potentially very distant bacterial lineages [10]. Transferred genetic material can correspond to single genes as well as clusters of consecutive genes on the chromosome from one or more transfer events. These latter are commonly described as genomic islands (GIs) [11]. They may bring an evolutionary advantage allowing adaptations to new environments or providing new pathogenicity capacities, for instance [12]. Studying the evolution and functional impact of GIs on bacterial populations is of major interest for microbiologists, since they are widely distributed in pathogenic and environmental microorganisms. GIs tend to insert at specific sites of the genome, such as tRNA genes [13]. Some of those insertion sites, called hotspots, are more active than the rest of the genome in terms of acquisition rate of new elements and have a much more diverse gene content, even between closely related individuals [14]. Indeed, hotspots diversify by rapid gene turnover driven by homologous recombination and horizontal gene transfer.

In the framework of the study of several *Escherichia coli* genomes, two publications have described the structure of GIs at a given hotspot [15, 16]. They showed a patchy structure corresponding to a segmented organization of genes into modules that can be found independently in different genome loci. These modules correspond to synteny blocks composed of consecutive genes that are conserved in a subset of the compared strains and can be functionally linked. In the *pheV* -tRNA hotspot of *E. coli*, the presence/absence of modules was shown to be uncorrelated with either the phylogenetic group or the pathotype [15]. Several other studies have described the modular structures of GIs in various species, such as in *Klebsiella pneumoniae* [17] or in *Photorhabdus/Xenorhabdus* [18]. Many methods to identify conserved synteny regions in prokaryotic genomes have been developed through the years. However, most of them do not scale higher than a few dozen genomes. To our knowledge, only two are designed for prokaryotes and have the ability to cope with more than a few hundred genomes, namely Gecko3 [19] and CSBFinder [20, 21]. While both of those methods provide an answer to the problem of finding conserved syntenies [22], they are not designed to work with pangenomes but rather on a more diverse taxonomic selection of genomes. Other approaches that do not rely on synteny conservation but on the cooccurrence or coevolution of genes in a pangenome to infer functional associations exist as well. Pantagruel [23] or Coinfinder [24] are recent examples designed for bacterial pangenomes. However, their goal is to infer associations or dissociations between pairs of genes, and they do not attempt to refine those relations into groups of colocalized genes directly.

In this article, we propose a new method called panModule to detect modules in GIs based on the pangenome graph representation of PPanGGOLiN [5]. In this graph, nodes represent gene families and edges represent genetic contiguity. In PPanGGOLiN, families are classified by a statistical model into a tri-partition scheme as introduced by [25]: (i) the persistent genome, which corresponds to genes that are present in most individuals of the studied clade, (ii) the shell genome, which groups genes that are conserved between some individuals of the group but not most and (iii) the cloud genome, which corresponds to genes that are rare within the population and found only in one or a few individuals. To predict modules, the panModule algorithm detects sets of co-occurring and colocalized genes in the variable part of the pangenome graph, composed of shell and cloud families. The predictions of panModule were evaluated at the species level using a curated dataset of modules in 12 genomes of *Escherichia coli* previously analyzed [15]. We then illustrate our approach by predicting modules in a set of complete *Klebsiella pneumoniae* genomes and comparing those predictions with formerly studied modules in a high pathogenicity island (HPI) of a strain of medical interest [17]. The panModule method is integrated in the PPanGGOLiN software suite (https://github.com/labgem/PPanGGOLiN). Modules can be predicted on pangenomes made of thousands of genomes and be analyzed along with the GIs and integration spot results from the previously published panRGP method [26].

## 2 Materials and Methods

### 2.1 Module detection algorithm

The panModule method uses the pangenome graph representation of PPanG-GOLiN [5] in which nodes correspond to homologous gene families (classified into *persistent, shell* and *cloud* partitions) and edges represent genetic contiguity (two families are linked in the graph if they contain genes that are neighbors in the genomes). Modules are defined as non-overlapping sets of cooccurring and colocalized variable gene families (*i.e., shell* and *cloud* families) that correspond to connected components in the pangenome graph. The graph algorithm behind panModule is inspired from a previously published method [27] that merges information from two or more graphs in a multigraph and then detects common connected components. These components correspond to conserved synteny groups, where the input graphs are genomes. In panModule, we do not use a multigraph representation but directly the pangenome graph to detect synteny conservation and thus modules in the variable genome.

First, each annotated genome sequence (*i.e*. complete genome sequences or contigs) is read to obtain a genome graph, which is a linear graph of genes that is cyclic in case of complete sequences of circular plasmids or chromosomes. Then, the pangenome graph *G*(*V, E*) is built from all genome graphs where *V* is a set of vertices corresponding to gene families and *E* is a set of edges representing genetic contiguity in the genome graphs between those gene families. An edge *e_i,j_* is added between each pair of gene families (*v_i_, v_j_*) if their corresponding genes are separated by less than *t* genes on a genome graph. For *t* ⩾ 1, it corresponds to a transitive closure applied to the genome graphs and allows to connect two families even if their genes are not directly adjacent. This can be useful in case a module in a genome is interrupted by a gene insertion (e.g. an insertion sequence) or has lost genes due to a deletion or pseudogenization event.

For each edge *e_i,j_*, two Jaccard similarity coefficients are computed as follows: 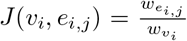 and 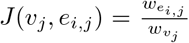 where *w_v_i__*. and *w_v_j__*. are the number of genes associated with the families *v_i_* and *v_j_*, respectively, and *w_e_i,j__* is the number of pairs of genes used to create an edge *e_i,j_* between two nodes *v_i_* and *v_j_*. A threshold *s* is defined as being the minimal Jaccard similarity to consider an edge as belonging to a module. If *J*(*v_i_, e_i,j_*) ⩾ *s* ∧ *J*(*v_j_, e_i,j_*) ⩾ *s*, the edge is kept, otherwise it is removed from the graph. In addition, nodes corresponding to gene families that are present in less than *m* genomes are also removed.

After this filtering step, each connected component is extracted using a modified Breadth-First Search (BFS) algorithm. A connected component is considered as a predicted module if it contains at least 3 nodes made of the *shell*, *cloud* or multigenic *persistent* families according to PPanGGOLiN classification [5]. Indeed, modules containing non-multigenic *persistent* families are not considered because they are not part of the variable genome and generally correspond to long syntenic regions that are conserved in almost all compared genomes without any or only a few rearrangement events.

### 2.2 Reference dataset and genome collections

To evaluate panModule predictions, we used a reference dataset of modules based on the expert annotation of 12 *E. coli* complete genomes originally published in [15]. These 12 strains are from different phylogroups (*i.e*. A, B1, B2 and D) and are commensal or ExPEC (Extra-intestinal Pathogenic *Escherichia coli*). Their genomes have been curated with the MicroScope platform [28] and their GIs have been divided into modules according to both synteny conservation and functional annotation of the genes. This dataset, named EcoliRef, contains a total of 165 modules that are present in at least 2, and not more than 10 of the 12 genomes, for a total of 793 occurrences in 461 GIs. GIs and modules are described by their genomic coordinates on the 12 *E. coli* chromosome sequences and were classified into 7 functional categories (Supplementary Data file ‘benchmark/reference_modules.tsv’).

Module prediction was run on 4 collections of *E. coli* genomes. The first one corresponds to the 12 genomes of the EcoliRef reference dataset. The second one contains all 1671 *E. coli* genomes classified as ‘Complete’ or ‘Chromosome’ in NCBI RefSeq [29] (downloaded the 1st of March 2021 and listed in Supplementary Data file ‘benchmark/EcoliComplete_genomes.list’). This dataset will be thereafter called EcoliComplete and includes the 12 genomes of EcoliRef. The third collection, named EcoliContigs, includes 1659 unfinished genomes plus the 12 genomes of EcoliRef. All *E. coli* genomes classified as ‘Contigs’ or ‘Scaffold’ in NCBI RefSeq (downloaded the 22nd of June 2021) were compared to all EcoliComplete genomes (apart from the 12 genomes of EcoliRef) with Mash (version 2.1.1, parameters: -s 5000, default for the others) [30] to iteratively pick out the closest genome in contigs (Supplementary Data file ‘benchmark/EcoliContigs_genomes.list’). As such, we get an equivalent bias in genome composition between both datasets to compare them properly and thus evaluate the impact of using genomes in contigs for module prediction. The last collection, named EcoliMAGs, contains 1 416 MAGs classified as ‘*Escherichia coli*’ from the study of Pasolli *et al*. [31] plus the 12 genomes of EcoliRef. MAGs were annotated using bakta (version 1.0.3, default parameters) [32]. As this dataset is much smaller than NCBI RefSeq, we chose not to apply the same filters as for the EcoliContigs dataset and just kept all MAGs. Therefore, we have here a collection of incomplete and fragmented *E. coli* genomes whose diversity is different from the two previous ones. Indeed, it contains potentially fewer pathogenic strains because MAGs were obtained from metagenomic samples not involving patients with an *E. coli* infections.

To illustrate the use of panModule on another species, a dataset containing 566 complete genomes classified as ‘s__Klebsiella pneumoniae’ in GTDB (release 06-RS202) [33] was downloaded on the 3rd of May 2021 from GenBank and NCBI RefSeq (listed in Supplementary Data file ‘Klebsiella_pneumoniae/Klebsiella_pneumoniae_genomes.list’). Then the modules were predicted using the method described in 2.1 with parameters *t* = 4, *m* = 2 and *s* = 0.86. The GIs of interest were identified by searching for those with positions overlapping with KPHPI208 in *Klebsiella pneumoniae* 1084. Annotations considered in the analysis are those from the downloaded file of NCBI RefSeq.

Genomic region illustrations were obtained using an online version of the CGView software [34] available at https://beta.proksee.ca.

### 2.3 Module prediction and benchmark procedure

The panModule method was run on the four *E. coli* genome collections using PPanGGOLiN software version 1.2.0 with default parameters to obtain homologous gene families and pangenome graphs. Module prediction was evaluated at the genome level using the reference modules from the 12 genomes of the EcoliRef dataset. In a GI of a given genome, a set of genes is considered to be part of a predicted module if their corresponding families are members of the same module. The benchmark uses genomic positions and processes GIs one by one with the following assumptions: (i) pairs of genes located between the genomic positions of a reference module are considered as positive relations (ii) pairs of genes that do not belong to the same reference module but are in the same GI are considered as negative relations. Thus, pairs of genes in a GI that are in the same predicted module and the same reference module are True Positives (TP). Pairs of genes that are in different predicted modules or not in a predicted module but in the same reference module are False Negatives (FN). Pairs of genes that are in the same predicted module but in different reference modules or not in a reference module are False Positives (FP). Pairs of genes that are in different predicted modules or not in a predicted module and in different reference modules or not in a reference module are True Negatives (TN).

To evaluate module prediction, Matthew’s correlation coefficient (MCC), F1-score, accuracy, precision and recall values were computed for the 4 *E. coli* genome collections as follows:

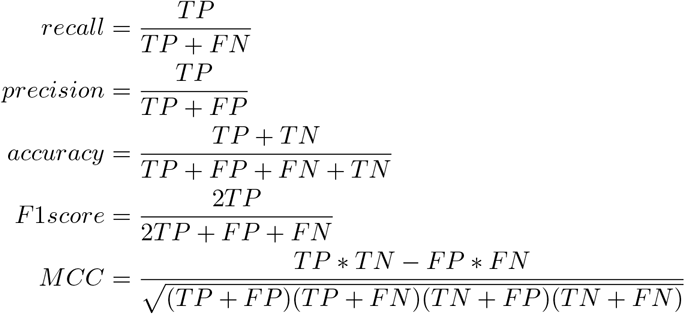

In order to determine the best values for the *s*, *m* and *t* parameters of panModule, we evaluated the predictions on the EcoliRef dataset using a set of realistic value combinations and chose the one that gives the best MCC (Supplementary Data file ‘benchmark/EcoliScope_benchmark_metrics.tsv’). We then used this set of parameters on the other genome collections for comparison.

## 3 Results & Discussion

### 3.1 Benchmark results

To evaluate panModule, we ran the benchmark as described in the Materials and Methods section to see how it performed against a curated set of functional modules. The best set of parameters was estimated on the EcoliRef dataset, and applied on the other datasets as such: *m* = 2, *t* = 4 and *s* = 0.86. A summary of the benchmark results for each dataset with these parameters is available in Table 1. It is possible that other parameters yield better results, which we will discuss with the case of the EcoliMAGs dataset.

**Table 1:** panModule results and benchmark on the four *E. coli* genome collections For each *E. coli* dataset, the number of genomes and pangenome gene families is given along with the number of families in the *shell*. Benchmark results are provided with two additional metrics (gene coverage and module coverage) corresponding to the percentage of genes and modules of the reference dataset that are associated to a predicted module. The benchmark was made using *s* = 0.86 except for the last column where *s* = 0.49. The number of predicted modules, their gene families and the percentage of shell families in modules are also provided.

| Metric | EcoliRef | EcoliComplete | EcoliContigs | EcoliMAGs | EcoliMAGs<br>($s = 0.49$) |
| --- | --- | --- | --- | --- | --- |
| Genomes | 12 | 1671 | 1671 | 1428 | 1428 |
| Families | 8867 | 57346 | 64318 | 44045 | 44045 |
| Shell families | 748 | 5542 | 5524 | 2410 | 2410 |
| MCC | 0.63 | 0.56 | 0.55 | 0.34 | 0.56 |
| F1 score | 0.66 | 0.59 | 0.60 | 0.31 | 0.60 |
| Accuracy | 0.85 | 0.83 | 0.83 | 0.76 | 0.83 |
| Precision | 0.96 | 0.93 | 0.89 | 0.90 | 0.90 |
| Recall | 0.51 | 0.43 | 0.45 | 0.19 | 0.45 |
| Gene coverage | 69.9% | 64.6% | 64.0% | 39.2% | 61.5% |
| Module coverage | 79.9% | 74.9% | 73.8% | 48.7% | 68.0% |
| Predicted modules | 219 | 1839 | 1516 | 1119 | 1485 |
| Families in modules | 1379 | 11558 | 9047 | 5473 | 11434 |
| Shell in modules | 61.9% | 48.0% | 42.2% | 33.2% | 54.8% |

The figure 1 represents the modules predicted with the different datasets on a plastic region of the genome of *Escherichia coli* 536 with the set of parameters previously mentioned.

**Figure 1:**
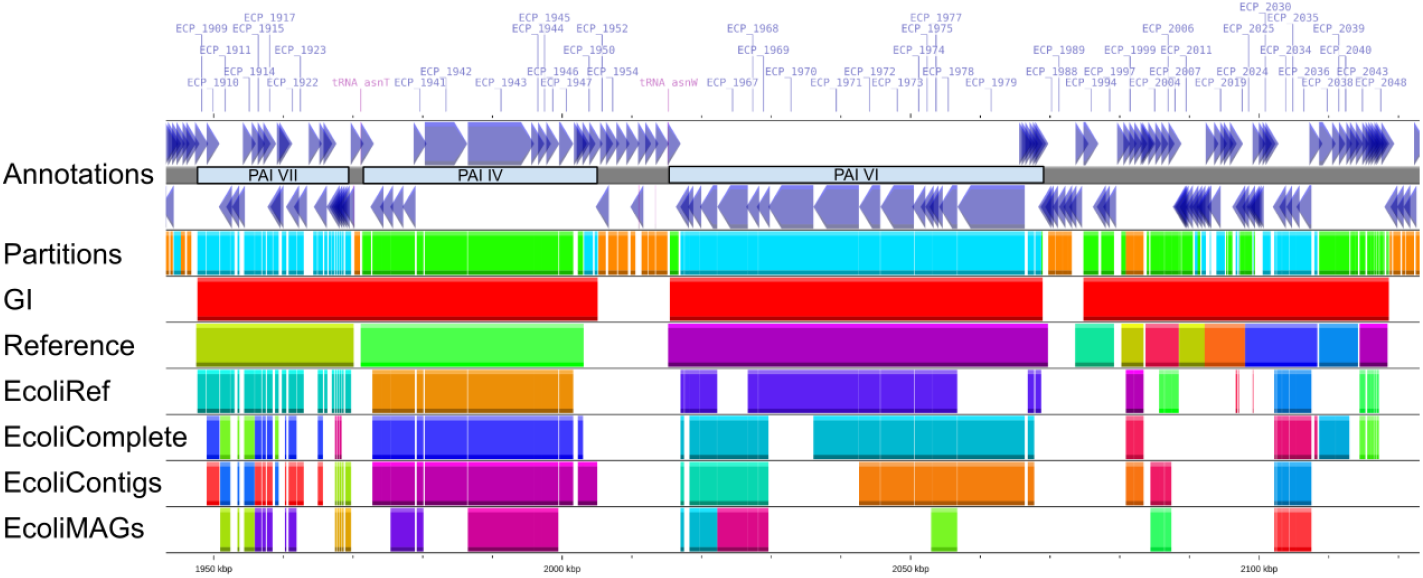
Modularized plastic region of *Escherichia coli* 536 Each track in the figure is a layer of information about genomic features in a region of the chromosome of *E. coli* 536 (accession CP000247.1). The 1st and 2nd tracks indicate gene positions and orientation. In between them are indicated the known pathogenicity islands in the region, respectively PAI-VII, PAI-IV and PAI-VI [35]. The 3rd track indicates for each gene its pangenome partition from the EcoliRef dataset by a color code: orange for *persistent*, green for *shell* and cyan for *cloud*. The 4th track indicates GI positions. The 5th track indicates reference modules. Tracks 6 to 9 show the module predictions using the different datasets (EcoliRef, EcoliComplete, EcoliContigs and EcoliMAGs). Colors for tracks 5 to 9 are selected randomly, and genes belonging to the same modules are colored identically in each track.

Overall, the modules detected by our approach in the EcoliRef dataset (*i.e*. the same set of genomes that were used for the expert annotation of modules) compare favorably to the reference modules with MCC and F1 score values of 0.63 and 0.66, respectively. The precision and accuracy values (0.96 and 0.85, respectively) are particularly high. This indicates that most of the genes belonging to a predicted module are found together in a reference module (True Positives) and that few genes are wrongly associated (False Positives). Similarly, most of the genes in a GI that do not belong to a same or any reference module are also not grouped into a module by our method (True Negatives). On the other hand, the recall value (0.51) is much lower, indicating that while we do recover proper modules a non-negligible number of the genes belonging to a reference module are not covered by our predictions (False Negatives). In summary, using the EcoliRef dataset, the modules predicted by panModule are frequently included in or equal to the reference modules and rarely overlap multiple reference modules, but some reference modules may be missed or be incomplete. Those missing genes represent about 30% of the genes associated to a reference module, and about 20% of the reference modules are not covered by any prediction (see Table 1 and Figure 1 as an illustration).

Both EcoliComplete and EcoliContigs datasets show equivalent results to the EcoliRef dataset but with slightly lower metrics overall, with the recall being the most impacted (*i.e*. 0.43 and 0.45, respectively, versus 0.51 for the EcoliRef dataset). It is further amplified when looking at the EcoliMAGs dataset whose metrics are much lower, especially in terms of recall, where it reaches a staggering value of 0.19, meaning that most of the reference modules are not predicted. Figure 1 clearly displays the sparsity of predictions in that specific dataset. However, it still keeps an acceptable precision and accuracy with 0.90 and 0.76, respectively, indicating that predicted modules even with the most incomplete and fragmented genome datasets are often actual modules. Indeed, the method parameters estimated on the EcoliRef dataset are likely too stringent to analyze MAGs as many modules may be incomplete or split on several contigs. We looked for the Jaccard similarity (*s*) parameter providing the best MCC using the same values for *t* and *m* (Supplementary Data file ‘benchmark/EcoliMAGs_benchmark_metrics.tsv’). The best value of *s* is 0.49, and gives a MCC value of 0.56 which is equivalent to the EcoliComplete and EcoliContigs datasets. Even using such a low *s* value, accuracy and precision remain high. Those results indicate that our method is applicable on very incomplete and fragmented genome datasets such as MAGs using a relaxed Jaccard similarity threshold.

This benchmark shows that panModule is able to correctly predict modules with a very good accuracy and precision even with incomplete genomes. It should be noticed that this validation is based only on a limited subset of curated data made of 12 *E. coli* genomes of which 6 are from the B2 phylogroup. Therefore, this reference dataset does not capture the overall diversity of *E. coli*. Nevertheless, panModule can modularize a large fraction of the *shell* pangenome of large and diverse genome datasets as well (e.g. 48% of the shell families are predicted in a module for the EcoliComplete dataset). It would have been interesting to validate the method on a larger amount of reference data and also on other species but we did not find such available resources.

### 3.2 Analysis of the KPHPI208 genomic island in *Klebsiella pneumoniae*

To illustrate the potential of panModule on other species, we chose to reanalyze the KPHPI208 GI of *Klebsiella pneumoniae* 1084. This 208-kb GI inserted at the asn-tRNA loci was described to be composed of 8 genomic modules (GM1 to GM8) using comparative genomics [17].

First, we used panRGP to predict GIs in *K. pneumoniae* 1084 from a pangenome made of 566 genomes. Two predicted GIs correspond to KPHPI208. The first detected GI starts at 1,744,478 bp and stops at 1,906,798 bp while the second one starts at 1,920,068 bp and stops at 1,952,189 bp. The two GIs cover the majority of the region, with the remaining portion being persistent genes between the two islands. In the KPHPI208 region made of 135 genes, panModule predicted 9 modules including a total of 98 genes. An overall picture of the region is represented in Figure 2 and Table 2 summarizes the correspondences between the formerly published and the predicted modules. The genomic positions of each described GMs and predicted modules are given in Supplementary Data file ‘Klebsiella_pneumoniae/Klebsiella_modules.tsv’.

**Figure 2:**
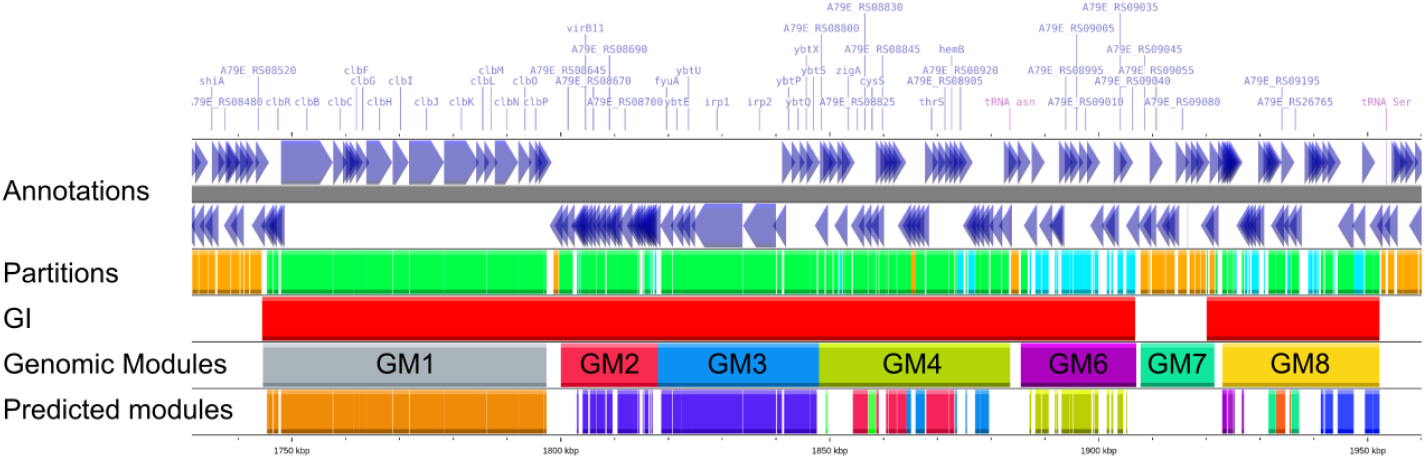
KPHPI208 genomic island in *Klebsiella pneumoniae* 1084 Each track in the figure is a layer of information about genomic features of the KPHPI208 genomic island. The 1st and 2nd tracks indicate gene positions and orientation. The 3rd track indicates for each gene its pangenome partition by a color code: orange for *persistent*, green for *shell* and cyan for *cloud*. The 4th track indicates GI predicted by panRGP. The 5th track indicates Genomic Modules (named GM1 to GM8). The 6th track indicates modules predicted by panModule.

**Table 2:**
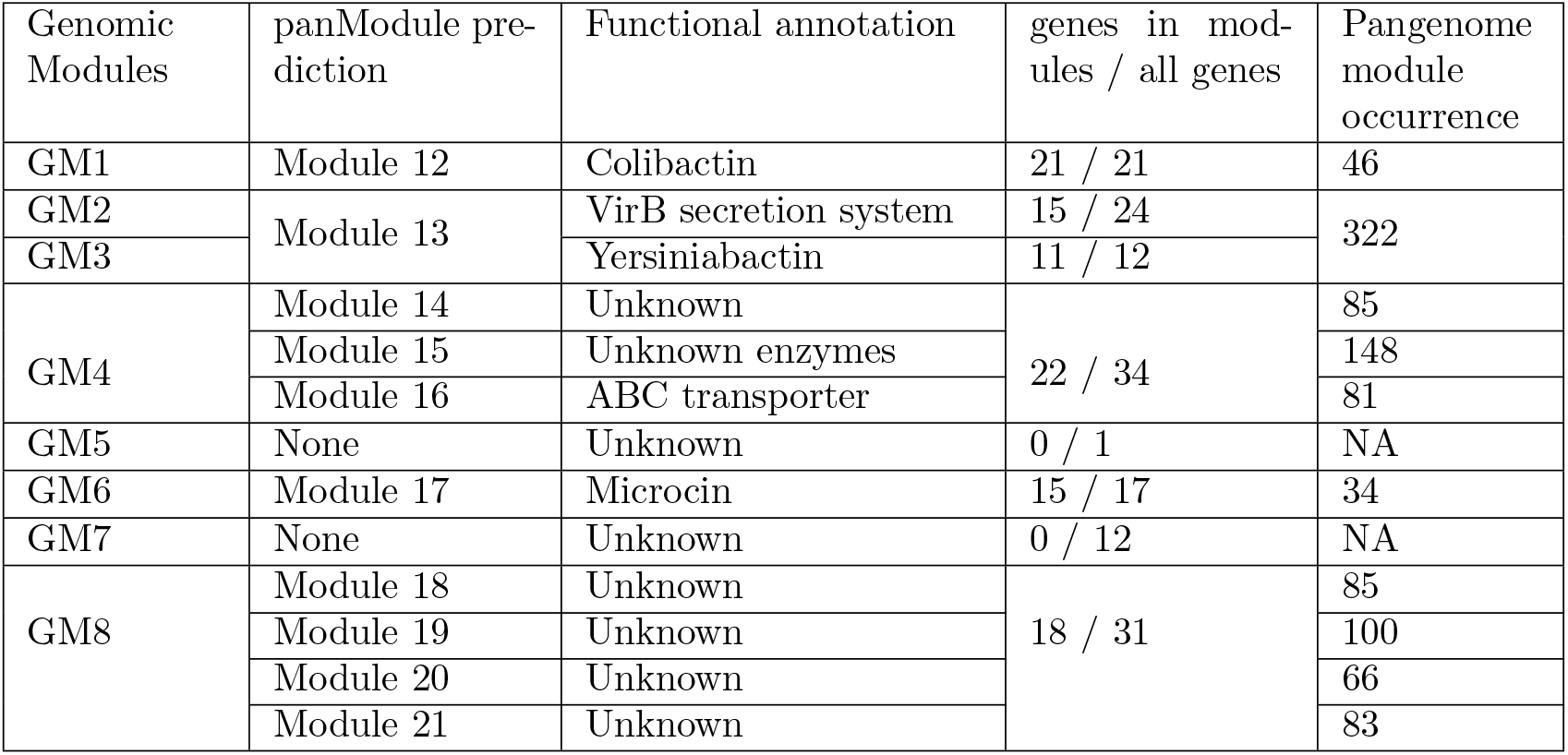
Comparison between genomic modules of the KPHPI208 genomic island and panModule predictions Genomic modules (GM) corresponds to the modules that were characterized in the original publication. The panModule prediction column contains the correspondences between the predicted modules and their GMs. Functional annotation column provides a brief summary of the module functions based on the gene annotation. Genes in modules / all genes column indicates how many genes are classified in a predicted module among all the genes in the original GM. Pangenome module occurrence column gives the number of genomes with the predicted module in the pangenome.

| Genomic Modules | panModule prediction | Functional annotation | genes in modules / all genes | Pangenome module occurrence |
| --- | --- | --- | --- | --- |
| GM1 | Module 12 | Colibactin | 21 / 21 | 46 |
| GM2 | Module 13 | VirB secretion system | 15 / 24 | 322 |
| GM3 |  | Yersiniabactin | 11 / 12 |  |
| GM4 | Module 14 | Unknown | 22 / 34 | 85 |
|  | Module 15 | Unknown enzymes |  | 148 |
|  | Module 16 | ABC transporter |  | 81 |
| GM5 | None | Unknown | 0 / 1 | NA |
| GM6 | Module 17 | Microcin | 15 / 17 | 34 |
| GM7 | None | Unknown | 0 / 12 | NA |
| GM8 | Module 18 | Unknown | 18 / 31 | 85 |
|  | Module 19 | Unknown |  | 100 |
|  | Module 20 | Unknown |  | 66 |
|  | Module 21 | Unknown |  | 83 |

Overall, our approach confirms the GMs that were functionally described by the authors, and provides novel insights for those that were uncharacterized. Only GM5 and GM7 were not predicted by panModule since they are composed of persistent genes. The predicted Module 12 is perfectly identical to the GM1 which codes for colibactin. Module 17 is extremely similar to GM6, only 2 genes which are likely unrelated to microcin biosynthesis were excluded from panModule predictions. Finding both GM2 (VirB) and GM3 (yersiniabactin) in the same predicted Module 13 indicates that they are conserved together in most *K. pneumoniae* genomes and suggests that they were exchanged together in the evolution of this species. Only two genes involved in the VirB secretion system are absent from the predicted module because they are not found conserved with other VirB system genes in a large number of genomes.

Among GMs that were described as unknown, GM4 and GM8 are composed of multiple modules predicted by panModule. Regarding GM4, we noticed that the genes that are in the same modules have similar annotations. No clear function could be inferred from Module 14. However, Module 15 contains mainly genes encoding enzymes and as such may be a metabolic pathway whereas Module 16 gathers proteins that are mostly related to an ABC transporter. Those common annotations are a strong indication that both are proper functional modules. For GM8, its analysis was less straightforward because most of the gene functions are either unknown or very generic and do not appear to be related to each other. Although we do not have much information in terms of functional annotation, the detected modules can be used as a basis for experimental studies to determine the biological processes in which they are involved.

Looking more broadly at the pangenome level, a total of 315 out of 566 *K. pneumoniae* genomes have a predicted variable region in the same integration spot as the one of GM1-GM6 region of the 1084 strain. It is a highly variable region, as there are 128 different organizations among the 315 genomes, with 108 different combinations of modules. The KPHPI208 organization appears only once in the pangenome as such, but 13 other genomes include all of those modules as well. A dynamic visualization of those 128 organizations with their module composition was generated by PPanGGOLiN and is available in Supplementary Data file ‘Klebsiella_pneumoniae/spot_19.html’.

## 4 Conclusion

We presented a novel method, named panModule, which groups genes into modules among thousands of genomes. Our approach uses a partitioned pangenome graph, which makes large-scale comparisons easier to compute. We benchmarked it against a curated set of *E. coli* genomes which were expertly annotated and whose GIs were divided into modules. We showed that panModule predictions were quite reliable regarding those annotations even for incomplete genomes. We illustrated the usefulness of our approach by revisiting the curated annotation of a genomic island in *K. pneumoniae* 1084. Overall, we believe that panModule provides an original approach to identify conserved modules in the variable regions of genomes, which may help to determine their function but also to better understand their complex evolutionary history.

The panModule method is freely available and easily installable as part of the PPanGGOLiN software suite, and can therefore be coupled with the other tools provided by the software, such as the analysis of pangenome partitions, the detection of GIs and spots of insertion. A potential improvement of the method presented here could be to include information from the species phylogeny in the computation of modules. Indeed, the calculation of Jaccard similarity could be weighted by a phylogenetic distance to favor the grouping of genes from distant strains into modules.

## 5 Data Availability

All mentioned Supplementary Data files and scripts to compute the benchmark are available at https://github.com/axbazin/panmodule-supplementary. The software is available at https://github.com/labgem/PPanGGOLiN.

